# Correction of the cytosine deamination artifacts in FFPE-based sequencing experiments

**DOI:** 10.64898/2026.08.11.744151

**Authors:** Wiktoria Płonka, Daria Kostka, Anna Lalik, Monika Kurpas, Khanh N Dinh, Magdalena Sitkiewicz, Marek Kimmel, Witold Rzyman, Roman Jaksik

## Abstract

Formalin-fixed, paraffin-embedded (FFPE) tissues remain an essential resource for molecular studies, yet formalin-induced cytosine deamination introduces characteristic C>T/G>A artifacts that compromise the accuracy of next-generation sequencing (NGS) analyses. Numerous computational methods and enzymatic DNA repair strategies have been proposed to reduce these artifacts, but no systematic comparison across tools and experimental conditions exists. Here, we evaluate the performance of seven computational approaches (SOBDetector, Ideafix, MicroSEC, FFPolish, DeepOmics FFPE/FFPE-PLUS, FFPErase) together with the NEBNext® FFPE DNA Repair Mix v2, a multi-enzyme repair system applied during DNA preparation. Using three independent datasets, one based on whole genome sequencing (CGCI-BL) and two on whole exome sequencing (TCGA-PC and SUT-LUAD, the latter containing enzymatically repaired samples), and matched fresh-frozen samples as the gold standard, we assess precision, sensitivity, and artifact reduction efficiency across all methods. We further examine the potential synergy between enzymatic repair and post-sequencing computational filtering. Our results provide practical guidelines for FFPE artifact correction and demonstrate that enzymatic treatment provides the best results, while among the computational methods, FFPErase offers the most robust reduction of cytosine deamination artifacts while maximizing the retention of true somatic variants.

**KEY MESSAGES:**

- Formalin fixation in FFPE samples introduces artifacts that can significantly affect the accuracy of NGS analyses.
- Among the evaluated approaches, enzymatic repair using NEBNext® FFPE DNA Repair Mix v2 achieves the most effective reduction of sequencing artifacts.
- Computational methods vary in performance, with FFPErase showing the most robust balance between artifact removal and retention of true somatic variants.
- Combining enzymatic repair with computational filtering did not lead to consistent improvements in performance across datasets.

## INTRODUCTION

Formalin-fixed, paraffin-embedded (FFPE) samples play a crucial role in biomedical research and clinical diagnostics, particularly in histopathological and molecular studies (Dodani *et al*. 2022). The long-term storage of tissue samples in FFPE allows for the analysis of biological material even years after it has been collected (Hedegaard *et al*. 2014). This method is widely utilized in hospitals, clinics, and scientific research, making FFPE a valuable source of data, especially for genomic studies such as next-generation sequencing (NGS) (Xuan *et al*. 2013). However, the efficiency of DNA isolation from FFPE tissues is relatively low, and the DNA obtained is of poor quality (Einaga *et al*. 2017). This is due to the properties of formalin, which interacts with DNA and introduces chemical modifications leading to the formation of DNA-DNA, DNA-protein (including DNA-histones), and DNA-formaldehyde crosslinks. The formation of crosslinks leads to changes in the structure of DNA and weakens the hydrogen bonds between DNA strands (Do and Dobrovic 2015). This results in DNA denaturation, increased susceptibility to DNA strand breakage, and decreased PCR amplification efficiency (Viljoen *et al*. 2022). The quality of DNA isolated from FFPE tissues depends on the method of fixation, the formalin solution used, and the DNA isolation method (Einaga *et al*. 2017). However, regardless of the method used, DNA isolated from FFPE tissues contains an increased number of sequence artifacts resulting from the removal or chemical modification of bases (Salgkamis *et al*. 2024). The most commonly described DNA damage caused by formalin is cytosine deamination, which leads to C>T alterations, that are mistakenly interpreted as mutations during genomic analysis (Tellaetxe-Abete, Calvo, and Lawrie 2021).

During the FFPE process, a 4% formalin solution permeates the sample, induces hydrolytic deamination of cytosine, converting it to uracil, which results in the creation of U:G mismatches. Consequently, DNA polymerases incorporate adenine instead of uracil, generating C>T substitution artifacts in the sequencing data (Guo *et al*. 2022). Uracil damage has been identified as a key source of sequencing artifacts in FFPE DNA. Among these artifacts, transient C>T variants represent the most common type of single nucleotide variation (SNV) (Do and Dobrovic 2015).

The presence of these artifacts in FFPE samples poses a significant challenge in accurately detecting mutations using NGS data, particularly when identifying cancer-related mutations with low allele frequency (Diossy *et al*. 2021). FFPE samples are reported to exhibit 3-4 times more mutations than fresh DNA (Quach, Goodman, and Shibata 2004). If these artifacts are not properly eliminated, they can lead to false-positive findings, severely undermining the reliability of the data (Tellaetxe-Abete, Calvo, and Lawrie 2021).

In response to this issue, several strategies have been introduced to mitigate the effects of cytosine deamination artifacts. These include computational methods designed to detect and remove potential errors after NGS data pre-processing (Diossy *et al*. 2021) and enzymatic techniques employed after DNA extraction from FFPE samples to minimize the number of formalin-related damages (Chen L *et al*. 2023).

Recently, a range of computational tools aimed at correcting FFPE artifacts, particularly those caused by deamination-induced mutations, have emerged: SOBDetector (Diossy *et al*. 2021), Ideafix (Tellaetxe-Abete, Calvo, and Lawrie 2021), MicroSEC (Ikegami *et al*. 2021), FFPolish (Dodani *et al*. 2022), DeepOmics FFPE (Heo *et al*. 2024), DeepOmics FFPE-PLUS(Kim *et al*. 2026) and FFPErase (Domenico *et al*. 2025). The basic features of all tools were summarized in table 1. All tools rely on quantitative properties of sequencing reads that help distinguish real variants from artifacts. A key measure is variant allele frequency (VAF), which reflects the fraction of sequencing reads supporting a mutation; FFPE-induced artifacts usually appear at low frequencies compared to true somatic variants (Do and Dobrovic 2015). Another common property is read orientation or strand bias, where artifacts tend to occur more often in reads aligned in one orientation of the DNA double strand, unlike true mutations that are more evenly distributed (Diossy *et al*. 2021). Related to this, strand imbalance captures systematic overrepresentation of variant-supporting reads on one strand (Diossy *et al*. 2021). Finally, some methods assess the distribution of mismatches within reads, since FFPE damage often creates clusters of errors at specific positions (such as read ends), whereas true variants are more evenly supported across sequencing reads (Ikegami *et al*. 2021). All seven tools exploit these properties to different degrees: SOBDetector focuses primarily on strand orientation bias, MicroSEC emphasizes error clustering and distorted read distributions within sequencing reads, Ideafix integrates variant allele frequency, mutational bias, and sequence context through decision-tree–based learning, FFPolish applies logistic regression across multiple quality and read-level features including variant allele frequency and strand bias, FFPErase uses 33 features per single nucleotide variant and 29 features per indel capturing base, mapping, positional, and contextual information, and DeepOmics FFPE employs deep neural networks to integrate a broad range of sequencing and caller-specific features (Heo *et al*. 2024).

**Table 1:**
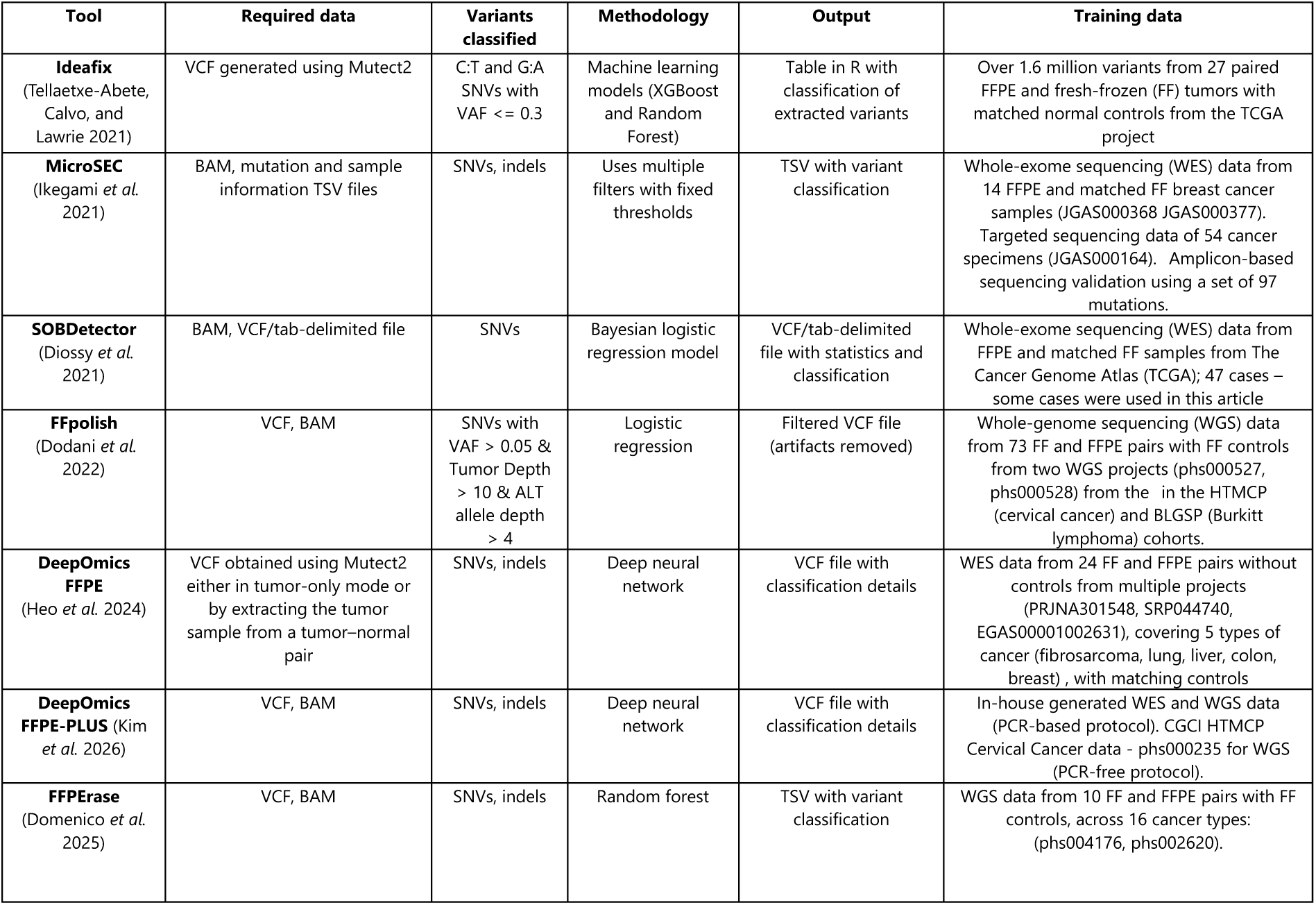
Existing computational artifact correction methods.

| Tool | Required data | Variants classified | Methodology | Output | Training data |
| --- | --- | --- | --- | --- | --- |
| **Ideafix**<br>(Tellaetxe-Abete, Calvo, and Lawrie 2021) | VCF generated using Mutect2 | C:T and G:A SNVs with VAF <= 0.3 | Machine learning models (XGBoost and Random Forest) | Table in R with classification of extracted variants | Over 1.6 million variants from 27 paired FFPE and fresh-frozen (FF) tumors with matched normal controls from the TCGA project |
| **MicroSEC**<br>(Ikegami *et al.* 2021) | BAM, mutation and sample information TSV files | SNVs, indels | Uses multiple filters with fixed thresholds | TSV with variant classification | Whole-exome sequencing (WES) data from 14 FFPE and matched FF breast cancer samples (JGAS000368 JGAS000377). Targeted sequencing data of 54 cancer specimens (JGAS000164). Amplicon-based sequencing validation using a set of 97 mutations. |
| **SOBDetector**<br>(Diossy *et al.* 2021) | BAM, VCF/tab-delimited file | SNVs | Bayesian logistic regression model | VCF/tab-delimited file with statistics and classification | Whole-exome sequencing (WES) data from FFPE and matched FF samples from The Cancer Genome Atlas (TCGA); 47 cases – some cases were used in this article |
| **FFpolish**<br>(Dodani *et al.* 2022) | VCF, BAM | SNVs with VAF > 0.05 & Tumor Depth > 10 & ALT allele depth > 4 | Logistic regression | Filtered VCF file (artifacts removed) | Whole-genome sequencing (WGS) data from 73 FF and FFPE pairs with FF controls from two WGS projects (phs000527, phs000528) from the in the HTMCPC (cervical cancer) and BLGSP (Burkitt lymphoma) cohorts. |
| **DeepOmics FFPE**<br>(Heo *et al.* 2024) | VCF obtained using Mutect2 either in tumor-only mode or by extracting the tumor sample from a tumor-normal pair | SNVs, indels | Deep neural network | VCF file with classification details | WES data from 24 FF and FFPE pairs without controls from multiple projects (PRJNA301548, SRP044740, EGAS00001002631), covering 5 types of cancer (fibrosarcoma, lung, liver, colon, breast) , with matching controls |
| **DeepOmics FFPE-PLUS** (Kim *et al.* 2026) | VCF, BAM | SNVs, indels | Deep neural network | VCF file with classification details | In-house generated WES and WGS data (PCR-based protocol). CGCI HTMCPC Cervical Cancer data - phs000235 for WGS (PCR-free protocol). |
| **FFPErase**<br>(Domenico *et al.* 2025) | VCF, BAM | SNVs, indels | Random forest | TSV with variant classification | WGS data from 10 FF and FFPE pairs with FF controls, across 16 cancer types: (phs004176, phs002620). |

Each tool was trained on a different dataset and evaluated using distinct benchmarking strategies, which makes direct comparison across methods challenging. SOBDetector was developed using FFPE sequencing data and validated by its ability to identify strand-biased artifacts compared to generic filters (Diossy *et al*. 2021). Ideafix was trained on over 1.6 million variants from matched FF and FFPE exomes and showed improved accuracy over conventional rule-based filtering when tested on independent datasets (Tellaetxe-Abete, Calvo, and Lawrie 2021). MicroSEC was assessed on 97 validated mutations, demonstrating high sensitivity and specificity in distinguishing true mutations from clustered sequencing errors (Ikegami *et al*. 2021). FFPolish used whole-genome datasets from Burkitt lymphoma and cervical cancer, and its logistic regression model was benchmarked against both single variant callers and combinatorial caller strategies, achieving higher precision and efficiency (Dodani *et al*. 2022). DeepOmics FFPE was trained on paired FF/FFPE whole exomes from multiple cancer types and comprehensively compared against SOBDetector, FFPolish, and Mutect’s FFPE filter; it removed 99.6% of artifacts while retaining more true variants, particularly at low allele frequencies, outperforming all compared tools (Heo *et al*. 2024). FFPErase is a random forest classifier trained using WGS data derived from multiple cancer types (Domenico *et al*. 2025). Most tools were compared against simple filtration methods however rarely to each other. MicroSEC and Ideafix were compared to SOBDetector in their respective articles while DeepOmics FFPE was compared to both FFPolish and SOBDetector, the most recent FFPErase was compared only to FFPolish. No comparison was made across all tools using a wide spectrum of data types, hindering the ability to systematically evaluate their relative performance, generalizability across cancer types, and robustness to different FFPE sample qualities.

An alternative to computational approaches are enzymatic methods of removing artifacts, which are used after DNA isolation from FFPE samples. These methods, rely on the use of specific enzymes that repair or eliminate damaged DNA fragments, potentially reducing the number of false positive variants (Chen L *et al*. 2023).

The most widely applied strategy targets cytosine deamination products. Uracil N-glycosylase (UNG) excises uracil from DNA, preventing the propagation of false C:G→T:A variants. Pretreatment of FFPE DNA with UNG reduces sequencing artifacts (Chen G *et al*. 2014, Do and Dobrovic 2015). Importantly, UNG was reported not to impair detection of true somatic variants (Chen G *et al*. 2014) and has been validated across diverse tissues and fixation times (Prentice *et al*. 2018, Berra *et al*. 2019). However, UNG treatment also has its drawbacks — the excision of uracil bases generates apurinic/apyrimidinic (AP) sites that are susceptible to cleavage under PCR conditions, which can elevate DNA fragmentation (Berra *et al*. 2019). Increased DNA fragmentation reduces the number of amplifiable DNA molecules, ultimately lowering library complexity and negatively affecting data yield in NGS experiments (Ottestad *et al*. 2022). For this reason, UNG is often integrated into commercial extraction kits (e.g., Qiagen QIAamp FFPE Tissue Kit) but is not always sufficient as a stand-alone solution.

Another approach involves the removal of single-stranded DNA regions through the treatment with S1 nuclease. Artifacts such as strand-split chimeric reads (SSARs) arise from single-stranded DNA fragments and overhangs. Treatment with S1 nuclease, which digests these single-stranded regions, removes those artifacts and thereby improves both single nucleotide variant (SNV) and copy number calling (Haile *et al*. 2019).

Recently enzymatic artifact correction has evolved from single-enzyme approaches targeting deamination to multi-enzyme BER-mimicking systems capable of addressing a wider spectrum of FFPE-induced DNA lesions. The NEBNext® FFPE DNA Repair Mix and its updated v2 (New England Biolabs) include glycosylases, endonucleases, polymerases, and ligases that collectively excise damaged bases, fill in gaps, and seal nicks. These treatments can reduce deamination and oxidation artifacts and partially restore sequence complexity (Chen L *et al*. 2023). In 2023 Steiert et al. introduced *In vitro* Sequential Base Excision repair (IQBErepair), a protocol using multiple glycosylases, including thymine-DNA glycosylase (TDG) and N-methylpurine-DNA glycosylase (MPG). Compared with NEBNext FFPE DNA Repair Mix (first version), IQBErepair achieved higher library complexity, more uniform coverage, and a stronger reduction of C>T and C>A artifacts (Steiert *et al*. 2023).

The major gap in the literature is the absence of studies that systematically compare all existing computational tools across multiple datasets and evaluate how their performance relates to enzymatic approaches. Another underexplored area is the potential synergistic use of both strategies - enzymatic and computational. A key question is whether combining these methods could provide better results than using either alone. In practice, such a hybrid approach could reduce artifacts more effectively, thereby improving NGS data quality and enabling more accurate research outcomes.

This study aimed to address these gaps by comprehensively evaluating the performance of seven computational methods for FFPE artifact removal in comparison with the latest commercially available enzymatic repair reagent kit - the NEBNext® FFPE DNA Repair Mix v2 Module, and the potential benefits of their combined application. For this purpose we utilized three distinct next generation sequencing datasets, one based on whole genome sequencing - CGCI-BL and two based on whole exome sequencing: TCGA-WES and our custom SUT-LUAD, which also includes the enzymatically repaired samples. By comparing FFPE samples with the corresponding fresh-frozen “gold standard” samples derived from the same tumors, we can assess the efficiency of various artifact detection methods. The experimental design is summarized in Figure 1.

**Figure 1:**
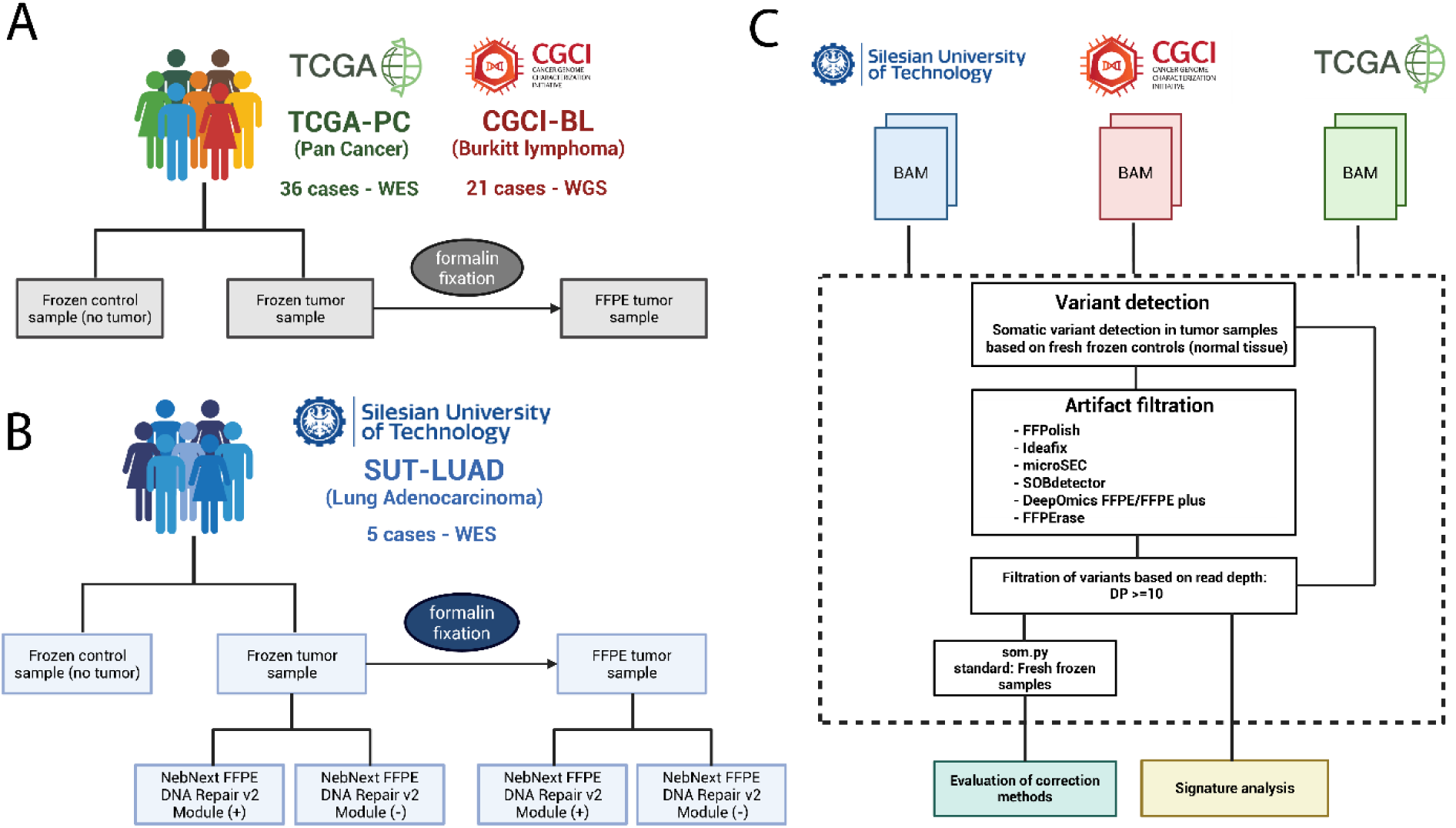
Experiment outline: A) Description of the TCGA-PC and CGCI-BL datasets, which include three different sample types obtained for a total of 57 cases; B) Description of the SUT-LUAD dataset, which comprises five sample types collected from five distinct LUAD individuals; C) Analysis outline, including variant detection, artifact filtration, and evaluation of filtration efficiency.

## MATERIALS AND METHODS

### Formalin treatment and enzymatic repair

The protocol was conducted on the primary tumor sample obtained from 5 lung adenocarcinoma patients (LUAD). Two pieces of similar size were cut from each tissue fragment examined. One piece was placed at −80°C (sample referred to as fresh frozen) and the other was fixed in neutral buffered formalin for at least 48h. DNA was isolated from each sample using the DNeasy Blood & Tissue Kit (QIAGEN) reagent kit according to the manufacturer’s recommended protocol. The isolated DNA, from both formalin fixed and fresh frozen samples, was divided into two parts and one part was treated with NEBNext® FFPE DNA Repair Mix v2 Module (New England Biolabs) according to the manufacturer’s protocol. All DNA samples ( treated and untreated with NEBNext® FFPE DNA Repair Mix) were then re-isolated using DNeasy Blood & Tissue Kit reagents.

### DNA sequencing

Whole exome sequencing was subcontracted to Macrogen Europe. Libraries were prepared using Agilent SureSelect V8-Post and sequenced using Illumina NovaseqX in 2×150bp mode to a minimum of ∼18Gbp/sample. The Q30 on all samples exceeded 94%.

### Data pre-processing and variant detection

Raw paired-end reads were subjected to adapter and quality trimming using Trim Galore (v0.6.10) with parameters *--length 20 --nextseq 20 --trim-n --paired*, removing low-quality bases and adapter contamination. Quality control of both raw and trimmed reads was performed using FastQC (Andrews 2010) (v0.12.1) and FastQ Screen (Wingett and Andrews 2018) (v0.15.2) to assess base quality, adapter content, and potential sample contamination.

Trimmed reads were aligned to the GRCh38 reference genome using BWA-MEM (Li and Durbin 2009, Li 2013) (v0.7.17) in alternative contig–aware mode. PCR duplicates were marked using the MarkDuplicates algorithm from the Picard toolkit (v2.18.26). Base quality score recalibration (BQSR) was performed with GATK (Auwera and O’Connor 2020) (v4.5.0.0) using known variant resources (dbSNP, Mills and 1000G gold standard indels, and known indels). The recalibrated BAM files were indexed and assessed for overall quality using SAMtools stats (Li *et al*. 2009)and FastQC. Coverage metrics were calculated with BEDTools (Quinlan and Hall 2010) (v2.31.1) relative to RefSeq transcripts and the Agilent SureSelect v8 exome capture regions.

Somatic variant calling was performed on tumor–normal sample pairs using GATK Mutect2 (v4.5.0.0). Variants were filtered using GATK’s FilterMutectCalls based on Mutect2 results, as well as sample contamination estimates obtained using CalculateContamination tool and read orientation bias statistics obtained with LearnReadOrientationModel tool. The filtered variant set was obtained using FilterMutectCalls, integrating both contamination estimates and read orientation bias metrics. All retained variants were functionally annotated using Ensembl Variant Effect Predictor (McLaren *et al*. 2016) (v107).

### Detection of sequencing artifacts

The detection of sequencing variants was conducted using the following tools:

- microSEC (v2.1.5) (Ikegami *et al*. 2021) For each sample, input TSV files required by microSEC were generated, including a mutation information file (containing mutation type, chromosome, position, reference and alternative allele, TRF repeat information, and neighborhood sequence) and a sample information file (sample name, mutation file path, BAM file, read length, adapter sequences, sample type, panel name, and BED file with repeat regions). For samples with a large number of variants, the analysis was performed separately for each chromosome. The output consisted of a TSV file classifying variants, from which those labeled as artifacts were filtered out from the original VCF files.
- Ideafix (v1.0.0) (Tellaetxe-Abete, Calvo, and Lawrie 2021) was run using the XGBoost classifier and the hg38 reference genome. Variants classified as potential deamination artifacts (“deamination” class) were removed from the VCF files. During the analysis, an issue was encountered with samples containing a small number of variants, where missing features (e.g., absence of variants downstream of a GC dinucleotide) prevented variable generation. To address this, the one_hot_encoding function was modified to allow execution even with an incomplete feature set.
- SOBDetector (v1.0.4) (Diossy *et al*. 2021) was executed using the original VCF and BAM files for each sample. The tool annotated variants with an artiStatus label in the VCF INFO field. Variants labeled as “artifact” were subsequently excluded from downstream analysis.
- FFPolish (v0.1) (Dodani *et al*. 2022) was executed using the reference genome and sample-specific BAM files. For each sample, the tool produced a filtered VCF file with FFPE-induced artifacts removed. Additional annotations were added to indicate whether variants were retained or removed during filtration.
- DeepOmics FFPE (Heo *et al*. 2024) analysis was conducted using the on-premise version of the tool, in collaboration with Theragen Bio. The same software is also accessible via the web portal at https://DeepOmics.co.kr/ffpe/home. In addition, the authors utilized DeepOmics FFPE-PLUS, a complementary algorithm designed specifically for the analysis of whole-genome sequencing (WGS) data, which is available at https://github.com/Theragen-Bio/DeepOmicsFFPE-PLUS. For the TCGA-PC dataset, technical filtering based on a minimum read depth (DP ≥ 10) was applied prior to algorithm execution to limit the query size. For both DeepOmics FFPE and DeepOmics FFPE-PLUS, variants were next filtered out based on the IS_VARIANT flag - only variants with IS_VARIANT != “artifact” (for DeepOmics FFPE) or IS_VARIANT != 1 (for DeepOmics FFPE-PLUS) were retained for downstream analysis.
- FFPErase (Domenico *et al*. 2025) was executed using the provided Nextflow script, which takes BAM and VCF files as input. The required mean insert size was obtained for each tumor BAM file individually using *samtools stats* (Li *et al*. 2009). Coverage statistics for WGS samples were calculated from the total number of aligned bases reported by *samtools stats*, while coverage for WES samples was computed as the mean coverage across capture regions using *bedtools coverage* (Quinlan and Hall 2010). The tool outputs a TSV file containing all annotated variants, including a column snvs_predicts that indicates which variants were classified as artifacts. Variants with TRUE in the snvs_predicts column were considered artifacts.

### Evaluation of artifact detection tools

Following artifact filtration, the resulting VCF files were compared to matched fresh-frozen (FF) samples using the som.py script from the hap.py package (Krusche *et al*. 2019) (v0.3.15), which enables accurate comparison of VCF files and reports key performance metrics, including sensitivity, precision, and the number of shared and unique variants. From the original VCF files, only variants explicitly classified as artifacts were removed; unclassified variants, or those not specifically labeled as artifacts, were retained for comparison but treated as a separate group. The specific procedure for generating the final VCF files for each tool, including artifact classification and filtering, is described in detail in the section Detection of sequencing artifacts.

To evaluate the performance of artifact filtration tools, we defined true positives (TP), false positives (FP), false negatives (FN), and true negatives (TN) based on comparisons between filtered FFPE and matched FF samples. Variants present in both FF and FFPE samples even after filtration were considered as TP. Variants absent from the FF sample but present in the FFPE sample (and not removed by the filtration process) were considered FP. Conversely, variants present in the FF sample but missing from the FFPE sample after filtering were considered FN. Finally, variants absent from the FF sample and correctly removed from the FFPE sample by the filtering tool were considered TN. All unclassified variants (omitted by SOBDetector and FFPolish) were treated as artifacts in the evaluation. This strategy is consistent with the filtering assumptions of each tool, which focus on removing low-confidence variants likely to represent technical artifacts, for which artifact classification might not be possible due to insufficient information. Ideafix is an exception since it classifies only C:T and G:A SNVs with VAF <= 0.3. In this case all unclassified variants were treated as true variants in the evaluation.

### Mutational signature and cosine similarity analysis

Single-nucleotide variants with DP ≥ 10) were selected for mutational signature analysis. Variants from fresh-frozen samples were used as a references, to compare the relative impact of each correction tool applied to individual FFPE samples.

Trinucleotide mutation matrices were constructed using the mut.to.sigs.input function from the *deconstructSigs* R package (Rosenthal *et al*. 2016) with the BSgenome.Hsapiens.UCSC.hg38 reference genome and normalized to relative frequencies by dividing each trinucleotide count by the total number of mutations per sample. Normalized mutational profiles were visualized in COSMIC format using the *sigminer* R package (Tao *et al*. 2023).

For each correction method, differences between FFPE and matched fresh-frozen profiles were calculated and displayed as symmetrical heatmaps. Similarity between FFPE and matched fresh-frozen profiles was quantified using cosine similarity. Statistical differences between uncorrected and corrected samples were evaluated using paired Wilcoxon signed-rank tests. All analyses were performed separately for the SUT-LUAD, TCGA Pan-Cancer, and CGCI-BL datasets.

## RESULTS

### Impact of formalin fixation on the detection of somatic SNVs

Across all three analyzed datasets: SUT-LUAD, TCGA-PC, and CGCI-BL we observed a consistent overrepresentation of variants with low variant allele frequency (VAF) in FFPE samples compared to their fresh-frozen counterparts (Fig. 2A). This suggests that formalin fixation introduces or preserves a significant number of low-frequency somatic SNVs, which may compromise the accuracy of downstream variant interpretation.

**Figure 2:**
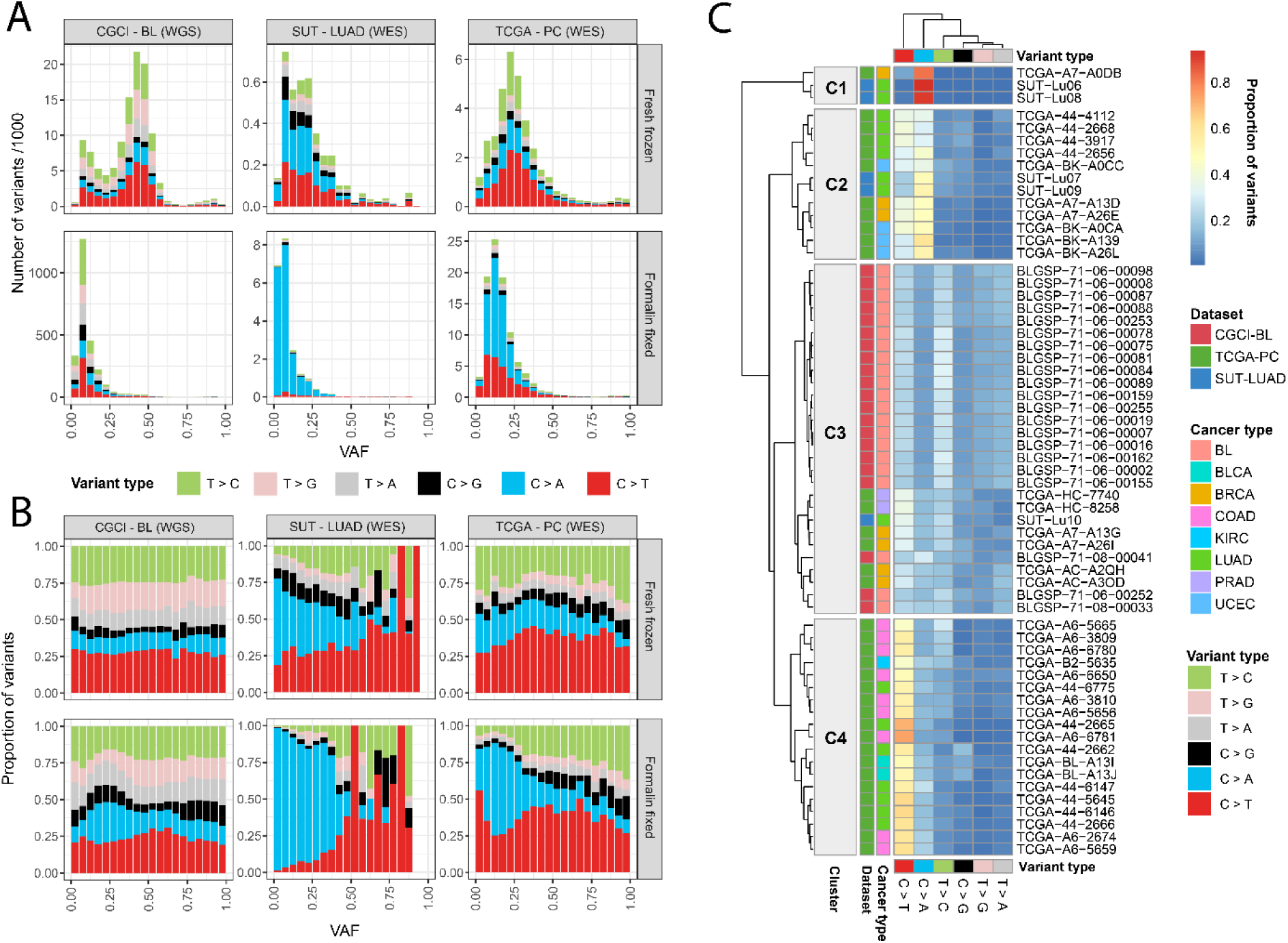
Variant characteristics identified in various datasets: A) Histograms of variant allele frequency (VAF) divided into specific variant types, prepared separately for each of the 3 datasets and Fresh Frozen and Formalin Fixed samples; B) proportion of variants corresponding to the plot A; C) heatmap of variant proportions for each individual sample

The distribution of specific substitution types varied markedly between datasets. In the SUT-LUAD dataset, C>A transversions dominated the FFPE variant spectrum (Fig. 2B), a mutational signature commonly associated with oxidative damage. In contrast, TCGA-PC samples showed a predominance of C>T transitions, with a secondary peak of C>A variants. The CGCI-BL dataset did not exhibit a clear preference for any single substitution type, but the total number of detected SNVs increased substantially post-fixation, further indicating a strong fixation-related impact on variant calling (Fig. 2A).

Clustering of samples based on mutational profiles revealed distinct groups with differing substitution patterns (Fig. 2C). Cluster C1, composed of three WES samples, was characterized by a strong overrepresentation of C>A mutations. Cluster C2, also including WES samples, displayed high levels of both C>T and C>A variants. Cluster C3 grouped all WGS samples along with selected samples from the other datasets and showed no dominant substitution pattern. Finally, Cluster C4, dominated by TCGA-PC samples, exhibited a clear excess of C>T transitions. Notably, these mutational biases were not correlated with cancer type, suggesting that the observed patterns are primarily driven by technical rather than biological factors.

### Correction of FFPE artifacts

The reported performance metrics are based on comparisons of filtered FFPE samples against matched fresh-frozen (FF) samples using *som.py* (Krusche *et al*. 2019). In the TCGA-PC (Pan Cancer) WES dataset, the highest precision and sensitivity values were achieved using FFPErase, FFPolish and DeepOmics FFPE. While the remaining algorithms demonstrated comparable sensitivity levels, their precision was substantially lower (Fig. 3A).

**Figure 3:**
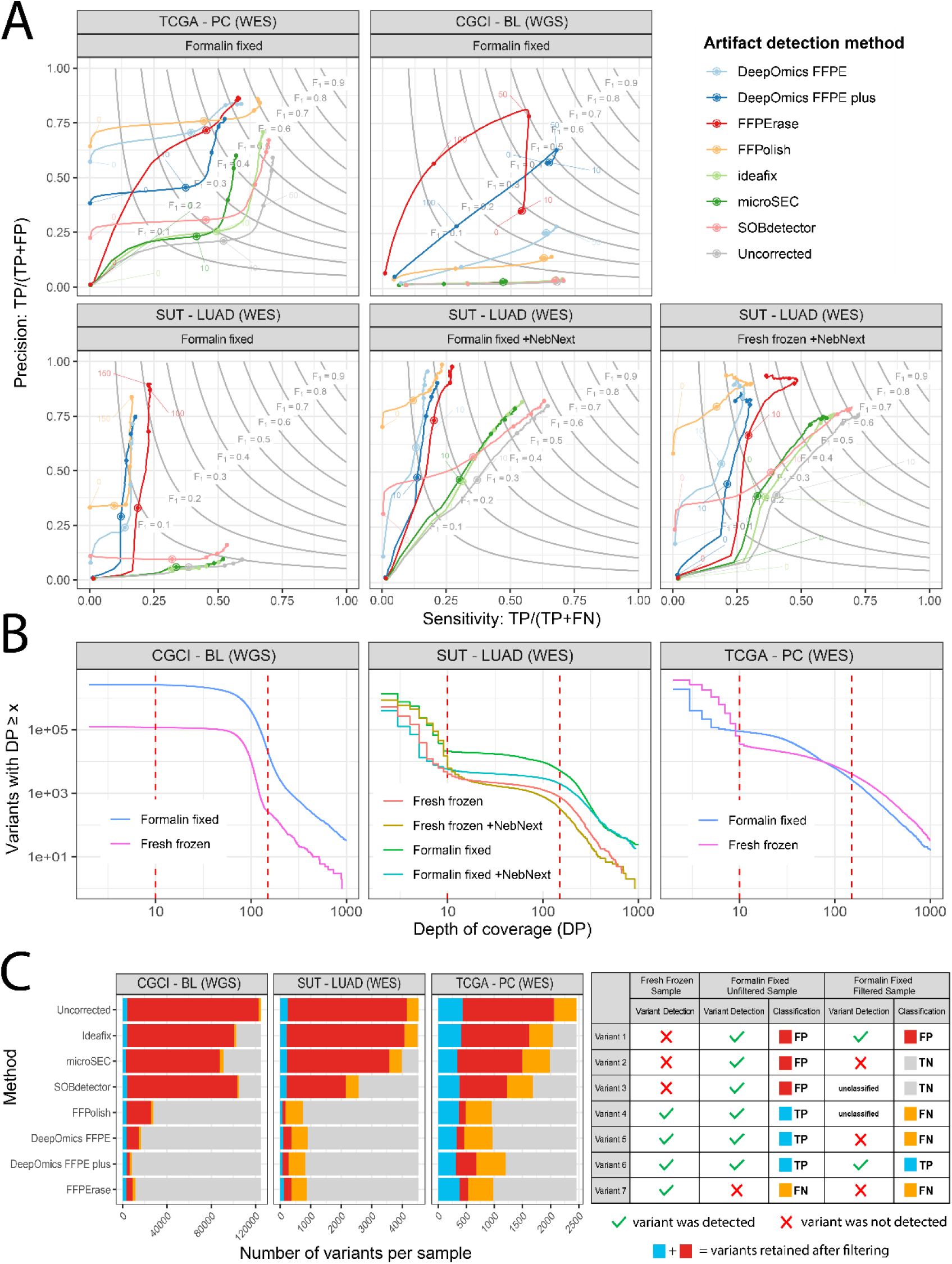
Accuracy of each of the variant filtration methods on Formalin Fixed samples compared to the Fresh Frozen: A) Sensitivity –precision curves calculated for variable coverage cutoff levels, gray lines mark the F1 score levels (harmonic mean of precision and sensitivity); B) Number of variants at specific coverage cutoff levels with 10x and 150x marked with red lines; C) Bar plots showing the average number of variants per sample divided into individual classes, assigned by each tool; TP - true positive, FP – false positive, TN – true negative, FN – false negative

For the CGCI-BL (WGS) dataset, the best performance was observed with DeepOmics FFPE-PLUS and FFPErase, both tools specifically designed for whole-genome sequencing data. Among the two DeepOmics FFPE-PLUS achieved the highest F1 score at 50x coverage cutoff level (DP ≥ 50) with only marginal decrease at 10x. FFPErase on the other hand performed poorly at 10x cutoff but showed significant improvement at 50x, and while it didn’t achieve higher F1 it significantly surpassed DeepOmics FFPE-PLUS at precision at the cost of sensitivity (Fig. 3A).

In the case of the SUT-LUAD (WES) dataset, the evaluated tools exhibited either high precision with very low sensitivity, or slightly improved sensitivity (by approximately 15% for variants with DP ≥ 10) accompanied by very low precision for formalin fixed samples. The highest F1 score was obtained using FFPErase, however none of the tested algorithms performed adequately on this dataset. However, when analyzing the same dataset but using FFPE samples prepared with the NEBNext FFPE DNA Repair protocol and followed by bioinformatic filtration, the results improved. Nevertheless, when compared to uncorrected samples, it appears that the improvement is primarily due to the enzymatic method rather than the filtration itself (Fig. 3A). However, it should be noted that all tools, especially FFPolish, tend to generate a considerable number of false negatives (FN), indicating the loss of true variants during filtering. This effect is particularly evident when comparing results to the Fresh Frozen + NEBNext FFPE DNA Repair samples.

Figure 3B illustrates the average number of variants per sample across each dataset as a function of minimum read depth thresholds. Vertical red lines indicate coverage cutoffs at 10× and 150×. WGS samples, as expected, show significantly more uniform coverage levels, with a higher number of variants retained even at more stringent DP thresholds, compared to WES. FFPE samples contain more variants than their matched FF counterparts, consistent with the presence of artifactual calls introduced during formalin fixation and processing. Importantly, the use of a DP ≥ 10 read depth filter effectively reduces the number of low confidence variants, improving the reliability of subsequent comparisons and performance evaluations. This filtering step is particularly useful in minimizing false positives in FFPE samples. Fig 3C additionally shows the number of variants per sample, stratified according to their classification into TP - true positive, FP – false positive, TN – true negative, FN – false negative. The plot shows the number of variants retained by each tool after filtering, which is the sum of TP and FP (blue and red).

### Impact of FFPE artifacts on single-base substitution frequency

Analysis of single-base substitution (SBS) patterns revealed notable differences between fresh frozen and formalin-fixed samples across most of the studied samples (Fig. 4A). In the SUT-LUAD cohort, most FFPE samples showed a significant increase in the fraction of C>A substitutions, with the strongest enrichment observed in the C[C>A]N trinucleotide context. This pattern is consistent with typical FFPE-induced oxidative damage and was similar across most individual SUT-LUAD samples.

**Figure 4:**
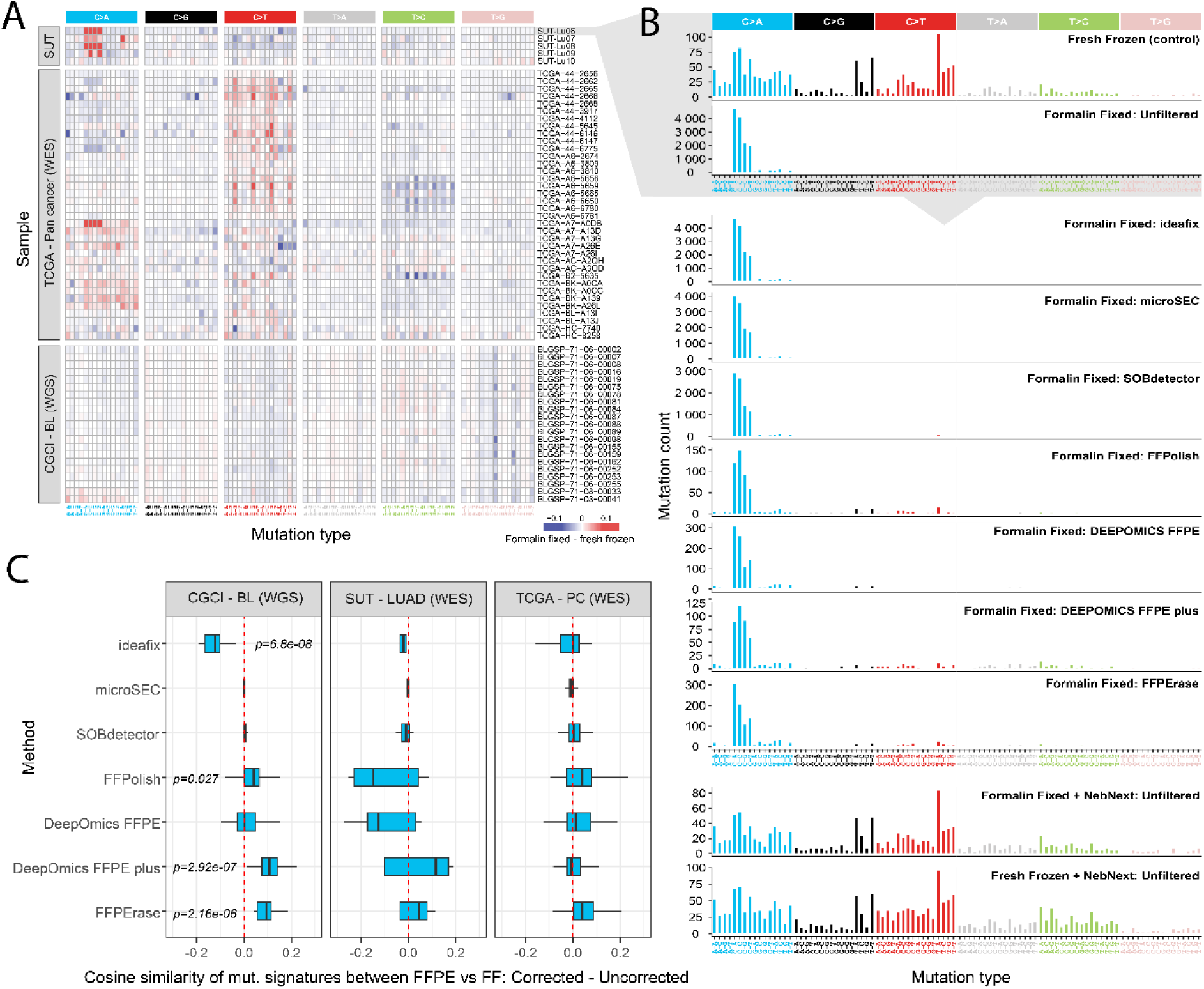
Single base substitution statistic for fresh frozen and formalin fixed samples before and after filtration: A) heatmap of the difference in variant proportion between fresh frozen and formalin fixed samples, for each individual patient; B) mutation counts divided into individual types for patient SUT-Lu06, the first two plots show the statistics used to calculate the difference in variant proportions shown on panel A. The six plots below show the signatures obtained after variant filtration using computational tools. Last two show the signatures obtained after treatment with NEBNext FFPE DNA Repair Kit; C) Boxplots of cosine similarity scores calculated between mutational signatures of fresh frozen and formalin fixed samples, shown as a difference between corrected and uncorrected samples; p-values originate from the Mann-Whitney test comparing the median difference relative to zero (red vertical line).

In contrast, the TCGA-PC dataset displayed a more heterogeneous picture. While an elevation in C>A substitutions was also observed in several TCGA-PC samples, only a single case exhibited a context-specific enrichment similar to the C[C>A]N pattern characteristic of the SUT-LUAD dataset. Instead, many TCGA-PC FFPE samples showed an increased fraction of C>T substitutions, predominantly in the N[C>T]G context, indicating the presence of additional context-dependent artifacts not shared across all datasets.

The CGCI-BL cohort, derived exclusively from unamplified WGS data, showed no clear FFPE-associated increase in the proportion of any specific mutation class. Instead, the dominant feature was an under-representation of T>G variants, most notably in the N[T>G]T context. This suggests that, in high-quality unamplified WGS data, FFPE artifacts may manifest differently or to a lesser degree than in the other datasets.

A detailed example is shown in Fig. 4B for sample SUT-Lu06. The first two panels represent the original signatures for the fresh frozen and FFPE materials, highlighting the strong FFPE-specific C[C>A]N excess. The subsequent panels illustrate the mutational spectra obtained after applying several computational correction strategies. All tested in silico filtering tools failed to fully restore the mutational profile to that of the matched fresh frozen sample. In contrast, enzymatic repair using the NEBNext FFPE DNA Repair Kit produced a substantially improved mutational signature, closely approximating the original fresh frozen profile. Importantly, treatment of a fresh frozen control with the same enzymatic protocol resulted in only minimal distortion of its baseline SBS distribution, confirming the specificity and limited off-target impact of the repair approach.

Cosine similarity scores quantifying the agreement between mutational signatures of matched fresh frozen and FFPE samples are summarized in Fig. 4C. Statistical comparisons using the Mann–Whitney test demonstrated that, while most computational methods failed to achieve a significant improvement over unfiltered FFPE data, several methods provided dataset-specific benefits. The highest similarity scores were obtained in the WGS-based CGCI-BL dataset after applying the DeepOmics FFPE-PLUS method, which, together with FFPolish and FFPErase, represented the only statistically significant enhancement relative to uncorrected data.

## DISCUSSION

The presented work shows that formalin fixation consistently increased the number of low-VAF calls and altered single-base substitution spectra, but the exact bias (C>T versus C>A enrichment and context specificity) varied between datasets and library protocols. This shows that FFPE artifacts are sample- and protocol-dependent and cannot be corrected with a universal rule.

Enzymatic repair using the NEBNext® FFPE DNA Repair Mix v2 produced the most significant and consistent improvement in mutational spectra and in agreement with matched fresh-frozen samples. In the SUT-LUAD cohort, enzymatic repair restored mutational profiles close to FF and substantially reduced FFPE-specific excesses that computational methods alone failed to remove.

Among computational methods, FFPErase provided the most robust improvement overall, when considering both WGS and WES data, achieving the best balance between artifact removal and preservation of true somatic variants. In the WGS dataset alone DeepOmics FFPE-PLUS showed the highest F1 score with FFPErase being only marginally worse at higher coverage cutoff levels. DeepOmics FFPE showed also relatively high performance in WES datasets, however it was outperformed by FFPErase. FFPolish performed well in WES datasets providing high precision, especially in the TCGA-PC WES data. The remaining tools (SOBDetector, Ideafix, MicroSEC) showed dataset-dependent performance: they can be useful in specific cases (e.g., where strand bias dominates), but their precision/sensitivity tradeoffs were inferior to DeepOmics FFPE in our benchmarks.

Enzymatic approaches, particularly single-enzyme UNG/UDG treatments are effective at removing uracil generated by cytosine deamination, but they are not free of drawbacks (Berra *et al*. 2019). Glycosylase activity creates AP sites, which can increase DNA fragmentation and reduce library complexity, especially when combined with mechanical shearing or other aggressive fragmentation steps (Steiert *et al*. 2023). This tradeoff can lower the number of amplifiable molecules and ultimately reduce sequencing yield and uniformity (Ottestad *et al*. 2022).

Multi-enzyme mixes, such as NEBNext® v2, attempt to mitigate these issues by coordinating excision and gap-filling steps. These approaches generally preserve library complexity better than single-enzyme workflows, though some loss of material and changes in coverage uniformity can still occur (McDonough *et al*. 2019, Chen L *et al*. 2023, Steiert *et al*. 2023). Laboratories should therefore validate enzymatic repair using representative sample types and extraction procedures, and should factor in the quantity and quality of the input DNA when selecting a repair strategy. It is also important to note that enzymatic repair reduces, but does not eliminate, all FFPE-associated artifacts. Residual lesions, including oxidative damage, crosslinks, and single-strand overhangs, may persist and can still benefit from computational filtering downstream (Haile *et al*. 2019).

Although our evaluation incorporated three independent datasets spanning both WGS and WES, and included enzymatically repaired samples (SUT-LUAD cohort), the broader diversity of fixation times, storage conditions, tissue types, and library preparation protocols found in real-world practice is greater than what we captured. As a result, performance may vary under conditions not represented here.

Several tools were originally developed and trained with specific variant callers and library protocols in mind. To ensure fairness, we used Mutect2 as a common caller because multiple evaluated methods expect Mutect2-originating VCFs. Using different upstream callers could shift absolute performance metrics, although the relative trends are likely to remain similar. Finally, although DeepOmics FFPE-PLUS performed best on WGS in our experiments, it is a relatively new tool and, like any machine-learning method, its performance will depend on how closely the target data resemble its training data. Some methods may be susceptible to implicit information leakage, if their training data share technical characteristics with the evaluation cohorts (e.g. sequencing chemistry, alignment and calling pipelines, DNA extraction protocols, or characteristic artifact signatures), performance may be partially inflated and not fully representative of generalization to independent data generated under different conditions.

Enzymatic repair of FFPE DNA (multi-enzyme BER-mimicking mixes) is the most effective single intervention for reducing cytosine deamination artifacts and restoring mutational spectra toward matched fresh-frozen profiles. Modern computational filters, particularly deep-learning methods such as DeepOmics FFPE/FFPE-PLUS — further improve data quality and are the recommended choice when enzymatic treatment is unavailable. Future work should expand validation across additional tissue types, fixation protocols and library preparations, and continue to evaluate emerging enzymatic protocols (including sequential BER strategies).

Practical recommendations regarding artifact filtration:

- When possible, perform enzymatic repair (multi-enzyme BER-mimicking mixes such as NEBNext® FFPE DNA Repair Mix v2) during sample preparation prior to library construction: this yields the greatest reduction in FFPE-specific artifacts and the only faithful reconstruction of mutational signatures. We however found this approach only applicable in cases where we had sufficiently large amounts of DNA available for a particular sample.
- If enzymatic treatment is not feasible, use FFPErase as a computational filter as the first choice, because it most consistently reduces deamination artifacts while preserving true low-VAF variants. For WES datasets DeepOmics FFPE and FFPolish are a reasonable alternative.
- In our datasets, hybrid strategies did not consistently improve results: enzymatic repair followed by a calibrated computational filter led in most cases to worse results (only SOBDetector showed a marginal improvement compared to unfiltered data), unless the experiment favors precision above sensitivity.
- Adopt conservative coverage and quality thresholds (we used DP ≥ 10 as a practical baseline) and inspect tool behavior across VAF bins and trinucleotide contexts before applying hard filters since artifact patterns are dataset-specific.
- Use Mutect2 for somatic calling in FFPE workflows: we used Mutect2 consistently because several evaluated tools were developed with Mutect2 output in mind and thus integrate most cleanly with its VCF annotations and flags.

## ACKNOWLEDGEMENTS

We would like to thank Inyoung Kim from Theragen Bio for conducting the DeepOmics and DeepOmics FFPE-PLUS analyses using the on-premise version of both tools.

The results published here are in part based upon data generated by the TCGA Research Network: https://www.cancer.gov/about-nci/organization/ccg/research/structural-genomics/tcga. and data generated by the Cancer Genome Characterization Initiative (CGCI; phs000235), *Burkitt Lymphoma Genome Sequencing Project (BLGSP; phs000527)*, developed by the National Cancer Institute (NCI). The data used for this analysis are available at the NCBI dbGaP (https://www.ncbi.nlm.nih.gov/projects/gap/cgi-bin/study.cgi?study_id=phs000527

Calculations were carried out using the infrastructure of the Ziemowit computer cluster (www.ziemowit.hpc.polsl.pl) in the Laboratory of Bioinformatics and Computational Biology, The Biotechnology, Bioengineering and Bioinformatics Centre Silesian BIO-FARMA, created in the POIG.02.01.00-00-166/08 and expanded in the POIG.02.03.01-00-040/13 projects.

## AUTHOR CONTRIBUTIONS

Wiktoria Płonka: Formal analysis (Supporting), Methodology (Supporting), Visualization (Supporting), Writing – original draft (Supporting). Daria Kostka: Formal analysis (Supporting), Visualization (Supporting), Writing – original draft (Supporting). Anna Lalik: Investigation (Lead), Writing – original draft (Supporting). Monika Kurpas: Formal analysis (Supporting), Methodology (Supporting), Project administration (Equal). Khanh Ngoc Dinh: Conceptualization (Supporting), Methodology (Supporting), Writing – review & editing (Supporting). Magdalena Sitkiewicz: Investigation (Supporting), Resources (Lead). Marek Kimmel: Conceptualization (Equal), Funding acquisition (Lead), Methodology (Supporting), Project administration (Equal), Supervision (Equal), Writing – review & editing (Supporting). Witold Rzyman: Resources (Equal), Supervision (Supporting). Roman Jaksik: Conceptualization (Lead), Data curation (Lead), Formal analysis (Lead), Methodology (Lead), Supervision (Equal), Visualization (Lead), Writing – original draft (Lead).

## SUPPLEMENTARY DATA

- **Supplementary_Tables_1.xlsx** – Tables associated with Fig.2

- SupplementaryTable 1.1: Number and fraction of variants at specific allele frequency (VAF) divided into substitution types, prepared separately for each of the 3 datasets and Fresh Frozen and Formalin Fixed samples
- SupplementaryTable 1.2: Variant proportions for each individual sample across all datasets SupplementaryTable 2.1: Variant filtration statistics calculated for variable coverage cutoff levels (minDP)
- **Supplementary_Tables_2.xlsx** – Tables associated with Fig.3

- SupplementaryTable 2.2: Number of variants at specific coverage cutoff levels
- SupplementaryTable 2.3: Average number of variants per sample divided into individual classes, assigned by each tool; TP - true positive, FP – false positive, TN – true negative, FN – false negative
- **Supplementary_Tables_3.xlsx** – Tables associated with Fig.4

- SupplementaryTable 3.1: Difference in single base substitution statistics between fresh frozen and formalin fixed samples, for each individual patient
- SupplementaryTable 3.2: Single base substitution statistics for all unfiltered and filtered samples
- SupplementaryTable 3.3: Cosine similarity scores calculated between mutational signatures of fresh frozen and formalin fixed samples, shown as a difference between corrected and uncorrected samples
- SupplementaryTable 3.4: p-values originating from the Mann-Whitney test comparing the median difference of cosine similarity score relative to zero

## DATA AVAILABILITY

Raw sequencing data from the SUT-LUAD dataset are available through the European Genome-phenome Archive (EGA) under the ID: EGAD50000001950.

## CODE AVAILABILITY

The complete code used in this study, including NGS data preprocessing, artifact detection, tool performance evaluation, and generation of all figures, is available on GitHub at: https://github.com/wplonka/Correction-of-the-cytosine-deamination-artifacts-in-FFPE

